# Autophagy Mitigates High-Temperature Injury in Pollen Development of *Arabidopsis thaliana*

**DOI:** 10.1101/088526

**Authors:** Gonul Dundar, Zhenhua Shao, Nahoko Higashitani, Mami Kikuta, Masanori Izumi, Atsushi Higashitani

**Author notes:** To whom correspondence should be addressed. 2-1-1, Katahira, Aoba-ku, Sendai, Miyagi 980-8577, Japan.

## Abstract

Autophagy is one of the cellular processes that break down cellular components during senescence, starvation, and stress. The susceptibility of plant pollen development to high-temperature (HT) stress is well known, but the involvement of autophagy in HT injury is yet to be clarified. Here, we found that following transfer to 30 °C, all autophagy-deficient (*atg*) mutants (*atg2-1, 5-1, 7-2*, and *10-1*) of *Arabidopsis thaliana* tested displayed visibly impaired pollen development and anther dehiscence. HT-induced male sterility significantly increased in the *atg* mutants, but the degree of HT-induced obstacles did not change between the wild type (WT) and mutants from the seedling stage to the bolting stage. Cytological analyses showed that 30 °C promoted autophagy and autolysosome formation in both anther wall cells and microspores in developing anthers of WT, but the *atg5-1* mutant did not show completion of tapetum degeneration and microspore maturation. HT upregulated hydrogen peroxide and dehydroascorbate reductase 1 production in both WT and *atg5-1* anthers, but the basal levels were already higher in the mutant. HT repressed expression of *UNDEAD* and its regulator *MYB80*, which are required for tapetal programmed cell death (PCD) for proper pollen development. Taken together, our results suggest that autophagy functions in tapetum degeneration and pollen development during HT-caused tapetal PCD abortion.

**Highlights:**

- In Arabidopsis, autophagy is not essential for completion of the life cycle under normal temperatures.
- High temperature (HT) stress induces autophagy in developing anther wall cells and microspores.
- Autophagy deficient *atg* mutants become almost completely male-sterile at moderate HT.
- Autophagy plays a role in tapetum degeneration and pollen development during HT-caused abortion of tapetal program cell death.

## 1. Introduction

Autophagy is a catabolic process that degrades cellular components within lysosomes and vacuoles, and is highly conserved among yeasts, plants, and mammals (Ohsumi 1999; Klionsky & Emr 2000; Yorimitsu & Klionsky 2005; Kim et al. 2007; Nakatogawa et al. 2009; Yoshimoto 2012). In plants, autophagy is activated during cell development, nutrient starvation, and senescence, and by environmental stressors such as pathogens, drought, salt, and oxidative damage (Bassham et al. 2006; Hofius et al. 2011; Yoshimoto 2012). In Arabidopsis and rice, a series of autophagy-related (*ATG*) genes have been identified, and many of the encoded essential amino acid residues are well conserved with those in yeasts (Doelling et al. 2002; Hanaoka et al. 2002; Chung et al. 2009; Xia et al. 2011; Yoshimoto 2012). Besides its role in cellular recycling, autophagy plays a role in cell death (Baehrecke 2005; Kroemer & Jäättelä 2005; Love et al. 2008; Parish & Li 2010). In plants, programmed cell death (PCD) occurs rapidly in the hypersensitivity response to pathogens, which requires prompt action to limit invasion, however autophagic cell death progresses relatively slowly and thus is observed in processes such as leaf senescence (Love et al. 2008; Parish & Li 2010; Wada et al. 2015).

Anther development, including specification of cell lineage and fate, follows well regulated programs. Among them, PCD is crucial in breaking down anther wall cells such as in the tapetum and middle layer during pollen grain maturation and anther dehiscence (Papini et al. 1999; Varnier et al. 2005). In Arabidopsis, rice, wheat, and cotton, the MYB80 transcription factor is required for the regulation of tapetal PCD through the expression of several genes, including *UNDEAD* (Phan et al. 2011, 2012; Xu et al. 2014). Autophagy plays a dispensable role in regular reproductive development, including tapetal PCD, in Arabidopsis, because almost all isolated autophagy-defective (*atg*) mutants can complete their life cycle under normal conditions (Yoshimoto 2012). Intriguingly, the occurrence of male sterility in a transgenic tobacco line overexpressing Arabidopsis *AtATG6/BECLIN1* in the tapetum cells (Singh et al. 2010, 2015) suggested that excessive autophagy might accelerate tapetum cell death and lead to abortion of microsporogenesis in tobacco. Nevertheless, the rice autophagy null mutant *OsATG7* showed a defect in tapetum cell degradation and sporophytic male sterility (Kurusu et al. 2014; Hanamata et al. 2014). These results strongly indicate that autophagy plays an important role in cell death during the degeneration of anther wall cells, but its biological significance and specific function in development are still unclear.

Early anther development is highly susceptible to several environmental stressors, in particular, high temperature (HT). We previously found that HT caused premature degradation of tapetum cells and abnormal vacuolization at 30 °C in barley and at >33 °C in Arabidopsis, and resulted in complete abortion of pollen development (Abiko et al. 2005; Oshino et al. 2007, 2011; Sakata et al. 2010). However, it is not clear whether elevated temperatures induce autophagy, and if so, how autophagy is involved in pollen and anther development under moderate HT stress. Here, we compared HT sensitivity during vegetative and reproductive stages between wild-type Columbia (Col-0; WT) plants and a series of *atg* mutants. We performed fine cytological analysis of anthers of WT and the *atg5-1* mutant, which is a typical *atg* mutant used in research on ATG8-dependent autophagosome formation (Thompson et al. 2005; Nakamura et al. 2018). We also analyzed gene expression and protein production following moderate HT of 30 °C.

## 2. Materials and Methods

### 2.1. Plant materials and growth conditions

We grew *A. thaliana* Col-0 and T-DNA knockout mutants *atg2-1* [SALK_076727], *atg5-1* [SAIL_129B07], *atg7-2* [GK-655B06], and *atg10-1* [SALK_084434], obtained from the Nottingham Arabidopsis Resource Centre, and recombinant plants of WT and *atg5-1* homozygously expressing the NahG, Salicylic acid (SA) hydroxylase (Yoshimoto et al. 2009), in chambers at 23 and 30 °C with an 8 h-light/16 h-dark photoperiod under fluorescent lamps (100 μmol m^−2^s^−1^). To test the effect of HT on seedling root growth, seeds were sterilized for 12 min in 5% (v/v) sodium hypochlorite containing 0.05% (v/v) Tween 20, washed with distilled water, kept at 4 °C for 3 days, and sown on 0.8% (w/v) Agar (Wako) containing 1/2 MS medium (Sigma-Aldrich) and 2% (w/v) sucrose in a plastic plate (ø 60 mm). Upon germination, plates were set vertically so that the seedlings grew straight along the surface of the medium. Seedlings with straight roots 1.0–1.5 cm in length were used.

To test the effect of HT on plant growth from seedling to bolting stages, seeds were directly sown into soil (1:1 mixture of Supermix A [Sakata] and vermiculite [Nittai]) and cultured at 23 or 30 °C in the growth chamber under the above conditions. To monitor the reproductive development and seed fertility of the primary inflorescence, 3-week-old plants (after initial anthesis) grown at 23 °C were transferred to 30 °C until the maturation of terminal siliques. To test recovery of pollen fertility, plants were held at 30 °C for 7 days, were then pollinated with pollen from plants of the same lines grown at 23 °C, and were kept at 30 °C.

### 2.2. *Construction of* ATG8e_pro_::YFP-ATG8e *recombinant plant*

To observe autophagosome formation, we generated an *ATG8e*-promoter–driven *YFP-ATG8e*–expressing plant (*ATG8e*_*pro*_::*YFP-ATG8e*). The protein-coding region of *ATG8e* (At2g45170) was amplified from Arabidopsis cDNA by reverse transcription (RT)-PCR with the ATG8e primer pair (ATG8e-F, 5′-CAC CAT GAA TAA AGG AAG CAT C; ATG8e-R, 5′-TTA GAT TGA AGA AGC ACC GAA TG), cloned into pENTR/D/TOPO (Invitrogen), and transferred to the pUBN-YFP vector (Grefen et al. 2010) to generate a *YFP-ATG8e* fusion. The DNA fragment containing this fusion was amplified using the YFP-attB1/B2R primer pair (YFP-attB1-F, 5′-GGG GAC AAG TTT GTA CAA AAA AGC AGG CTT TAT GGT GAG CAA GGG CGA GGA GCT G; ATG8e-attB2R-R, 5′-GGG GAC AGC TTT CTT GTA CAA AGT GGT TAG ATT GAA GAA GCA CCG AAT G) and cloned into pDONR221 (Invitrogen). The promoter region of *ATG8e*, which includes a 2607-bp upstream region from the start codon, was amplified from Arabidopsis genomic DNA using the ATG8ePro-attB4/B1R primer pair (ATG8ePro-attB4-F, 5′-GGG GAC AAC TTT GTA TAG AAA AGT TGA TCG CAC GGT CCC AAT ATG; ATG8ePro-attB1R-R, 5′-GGG GAC TGC TTT TTT GTA CAA ACT TGC TTT GCT TCT GAG AAT ATA CAC AAT C) and cloned into pDONR P4-P1R (Invitrogen). To generate the construct expressing the *YFP-ATG8e* fusion under the control of the *ATG8e* promoter, we transferred the pDONR221 plasmid containing the *YFP-ATG8e* fusion into the R4pGWB501 vector (Nakagawa et al. 2008) with the pDONR P4-P1R plasmid containing the promoter region of *ATG8e*. The resulting vector was transformed into Col-0 plants by *Rhizobium radiobacter* through the floral dip method (Clough & Bent 1998). Genotypic analysis of the segregating populations was performed in the T_2_ generation, and each T_3_ homozygous line was used for expression analysis.

### 2.3. Cytological analysis

To see HT-induced formation of YFP-ATG8 foci in anther wall cells and microspores, we observed developing anthers dissected from 3-week-old *YFP-ATG8e*-expressing plants grown at 23 °C and after 3 days’ treatment at 30 °C under a laser scanning confocal microscope. The number of YFP-ATG8e foci per anther cell was quantified by IMARIS Microscopy Analysis Software (Bitplane).

For transmission electron microscopy, dissected stage 9 anthers of plants grown at 23 °C and exposed to 30 °C for 3 days were fixed in 2% paraformaldehyde and 2% glutaraldehyde in 0.05 M cacodylate buffer (pH 7.4) at 4 °C for 24 h. They were then washed 3 times in 0.05 M cacodylate buffer for 30 min each, and fixed in 2% osmium tetroxide at 4 °C for 4 h. They were dehydrated through a 70% to 100% alcohol series for 30 min each, rinsed twice for 1 h in propylene oxide, and embedded in resin (Quetol 651; Nisshin EM Co.) for 48 h at 60 °C. Ultra-thin sections (80 nm) cut with a diamond knife on an ultramicrotome (Ultracut UCT, Leica) were mounted on copper grids, stained with 2% uranyl acetate at room temperature for 15 min, secondary-stained with lead stain solution (Sigma-Aldrich) at room temperature for 3 min, and then examined by transmission electron microscopy (JEM-1400 Plus, 80 kV; JEOL).

Transverse sections of stage 13 anthers and of pistils from 7-day HT-treated flowers at anthesis were prepared from embedded samples as above at a thickness of 1 µm and stained with 0.05% toluidine blue. Specimens were observed under a light microscope (BX51 Olympus) fitted with a CCD camera (DP73 Olympus). Dissected stage 13 anthers were also stained with iodine solution (Merck) to reveal pollen development and anther dehiscence.

### 2.4. DAB staining

Developing anthers were stained with 3,3′-diaminobenzidine (DAB: Sigma-Aldrich) as previously described (Orozco-Cardenas and Ryan 1999). Stage 12 anthers were dissected from unopened buds of Col-0 and *atg5-1* mutant plants either treated for 3 days at 30 °C or not, and infiltrated with DAB staining solution (1 mg mL^−1^ of DAB, pH 3.8) under vacuum. After 7 h incubation in darkness with gentle shaking, the anthers were washed and observed under a microscope. The intensity of staining was quantified in ImageJ software with 4 different samples per treatment.

### 2.5. *Quantitative analyses of* MYB80 *and* UNDEAD *genes*

Total RNA was isolated from 30 to 50 stamens at anther stage 9 in Trizol Reagent (Invitrogen). Real-time quantitative RT-PCR was performed with a PrimeScript II 1st strand cDNA Synthesis Kit (TaKaRa Bio) and SYBR Premix Ex Taq II (TaKaRa Bio), in a CFX96 Real-Time System (Bio-Rad Laboratories), with the following forward and reverse primers: 5′-GTT TCA CTC TGT TCT TGG TAA CC −3′ and 5′-CGG ATC TAT TCC CAT TCC TGA C-3′ for *MYB80*, 5′-ATG AAA ACC ACA ATG AAT TTT GTT TTT C-3′ and 5′-CTA CAT ATC ACA ATC TTG TTT ATT AAT-3′ for *UNDEAD*, and 5′-CCA GCT TTG GTG ATT TGA AC-3′ and 5′-CAA GCT TTC GGA GGT CAG AG-3′ for *Tubulin 2/3*.

### 2.6. *Semi-quantitative analyses of protein expression by* MALDI-TOF MS

A total of 20 µg of crude extract from 50 stage 12 anthers was separated by SDS PAGE in a precast 5%–20% gradient gel (Oriental Inst HOG-0520). Three independent gels were stained with Coomassie Brilliant Blue (Sigma-Aldrich). The intensity of each band was calculated in ImageJ software, and the major proteins in each band were determined by matrix-assisted laser desorption/ionization – time-of-flight mass spectrometry (MALDI-TOF MS; ABI5800, Applied Biosystems). Changes in relative levels of specific proteins by moderate HT treatment at 30 °C for 3 days was semi-quantified against specific peptide peak signals described in a previous protocol (Sasagawa et al. 2005): glutathione S-transferase / dehydroascorbate reductase 1 (DHAR1), L-ascorbate peroxidase 1 (APX1), and cytosolic triose-phosphate isomerase (CTIMC) were normalized to 60S ribosomal protein L7-2 (RPL7B) in MS spectrum data in the same gel slice from the 24–26-kDa protein band with the following peptides identified by MS/MS analysis: SHDGPFIAGER (theoretical molecular weight 1184.5) for DHAR1, ALLDDPVFRPLVEK (theoretical molecular weight 1610.9) for APX1, VIACVGETLEER (theoretical molecular weight 1374.6) for CTIMC, and ENFINELIR (theoretical molecular weight 1146.6) for RPL7B.

### 2.7 Statistical analyses

Each series of experiments was performed in triplicate. Statistics were calculated in MS Excel software. Statistical significance was assessed by unpaired Student’s two-tailed *t*-test. Values were considered statistically significant at *P* < 0.05 or *P* < 0.01. One-way ANOVA followed by Tukey’s *post hoc* test was performed for multi-sample study of silique length at *P* < 0.05.

## 3. Results

### 3.1. Autophagy-defective mutants showed hypersensitivity to HT stress during reproductive stage and resulted in male sterility

In *Arabidopsis thaliana*, >33 °C HT causes male sterility owing to the abortion of pollen development (Sakata et al. 2010). To study the contribution of autophagy to HT injury, we measured silique length and seed set in 3-week-old plants (after initial anthesis) of WT and *atg* mutants moved from 23 °C to moderate HT of 30 °C. In WT, fully elongated siliques from the 5th to the terminal blossoms were shortened to ½ to ⅓ of the length of those kept at 23 °C (Fig. 1A). In all *atg* mutants, they were reduced more severely to ⅙ of their length at 23 °C, which was normal (Fig. 1A). These shortened siliques in the *atg* mutants withered early. After the shift to 30 °C, the number of seeds per silique from the 5th to the terminal blossoms decreased to ¼ in WT and to almost nil in the *atg* mutants (Table 1). These results indicate that the 5th flower (flowering stage 10–11, Smyth et al. 1990; anther stage 9, Sanders et al. 1999) is a critical stage for susceptibility to elevated temperature, and that *atg* mutants became hypersensitive to HT stress during the reproductive stage.

**Table 1.**
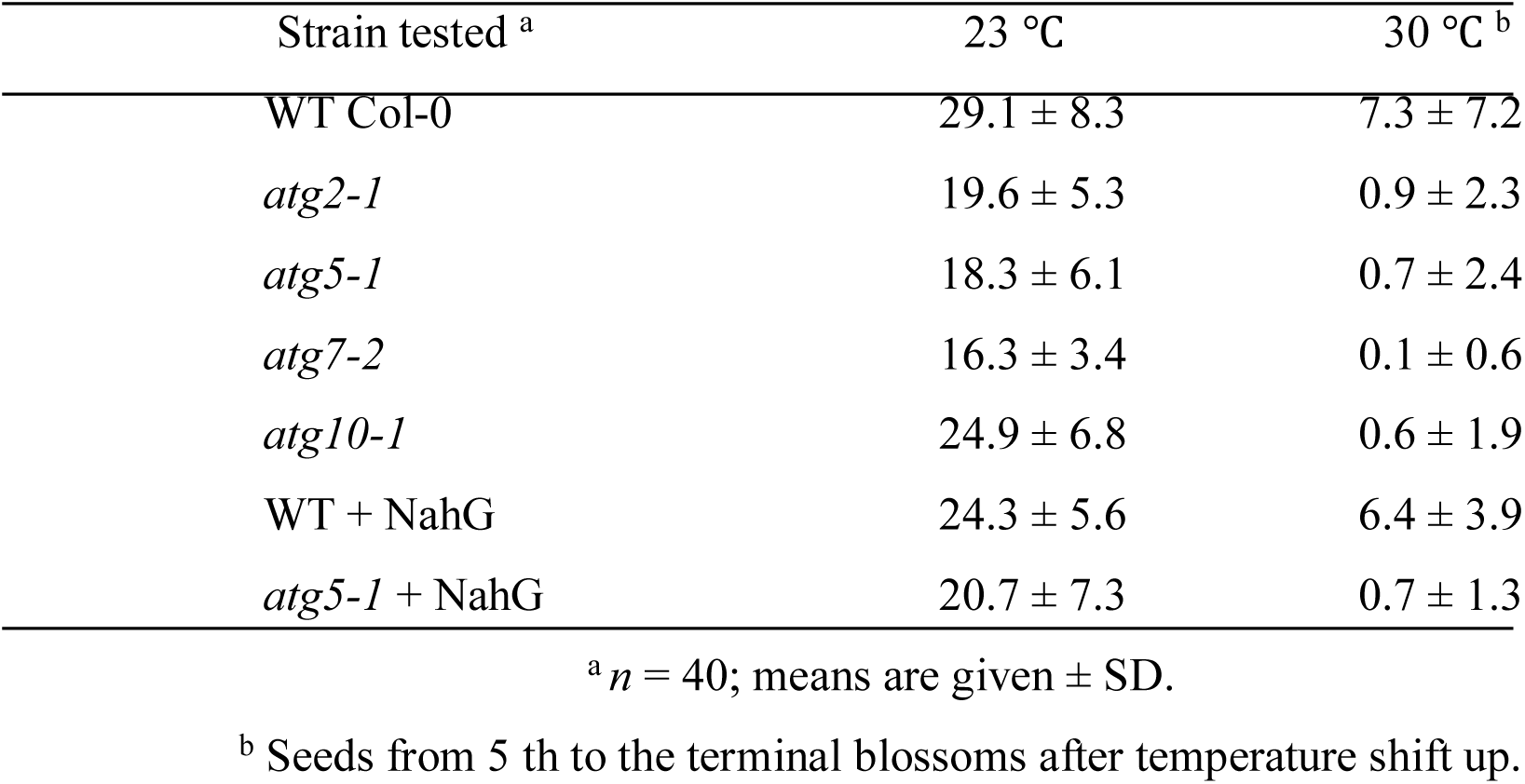
Number of mature seeds per silique in wild-type Col-0 and *atg* mutants

| Strain tested <sup>a</sup> | 23 °C | 30 °C <sup>b</sup> |
| --- | --- | --- |
| WT Col-0 | 29.1 ± 8.3 | 7.3 ± 7.2 |
| *atg2-1* | 19.6 ± 5.3 | 0.9 ± 2.3 |
| *atg5-1* | 18.3 ± 6.1 | 0.7 ± 2.4 |
| *atg7-2* | 16.3 ± 3.4 | 0.1 ± 0.6 |
| *atg10-1* | 24.9 ± 6.8 | 0.6 ± 1.9 |
| WT + NahG | 24.3 ± 5.6 | 6.4 ± 3.9 |
| *atg5-1* + NahG | 20.7 ± 7.3 | 0.7 ± 1.3 |

<sup>a</sup> *n* = 40; means are given ± SD.
<sup>b</sup> Seeds from 5<sup>th</sup> to the terminal blossoms after temperature shift up.

**Figure 1.**
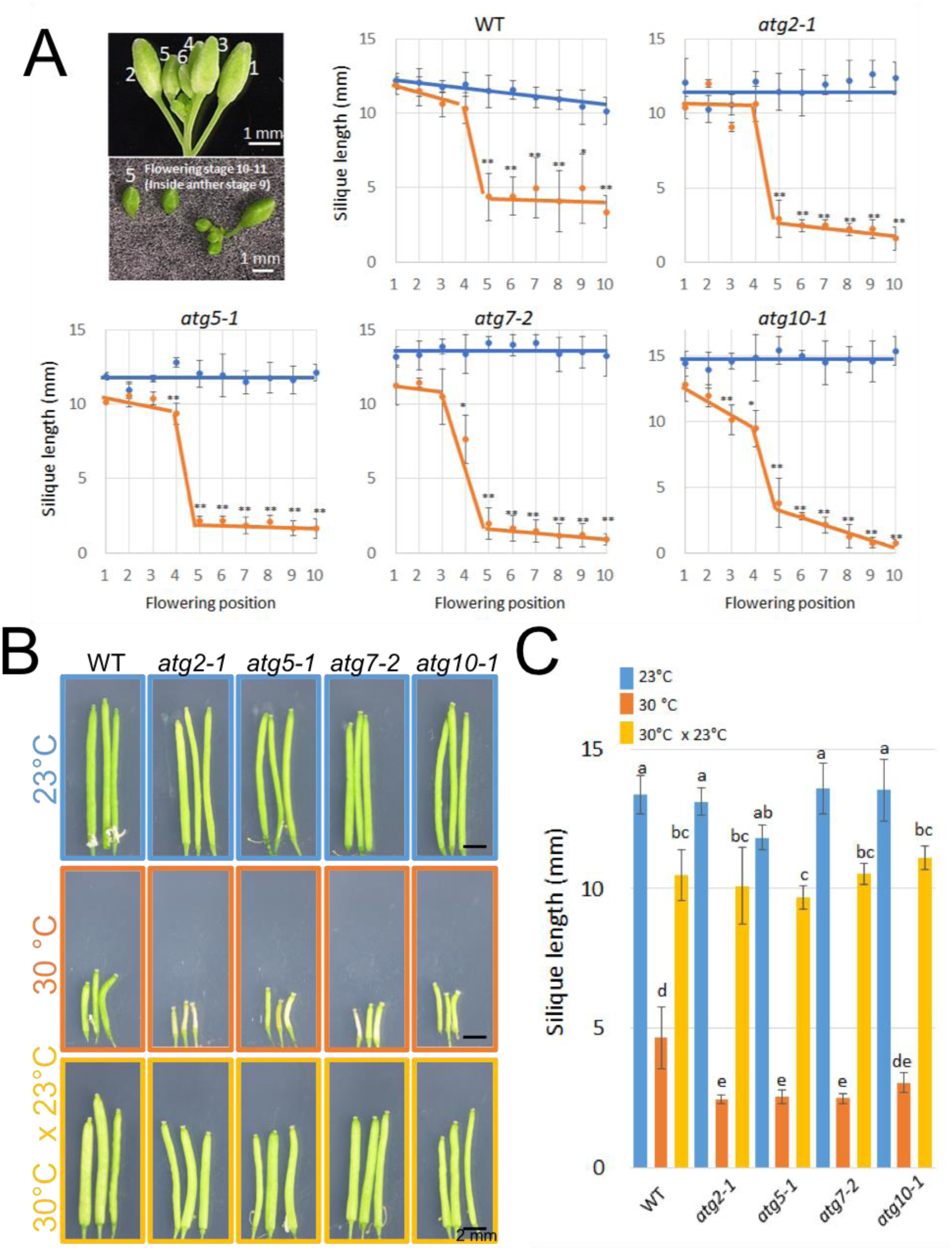
HT stress at 30 °C severely decreases silique length of *atg* mutants. (A) Silique length in each flowering position of primary inflorescence after transfer of 3-week-old plants of WT and *atg* mutants grown at 23 °C to 30 °C. At 23 °C (blue), silique length was unaffected. At 30 °C (orange), silique length was drastically reduced from the 5^th^ flower (identified in inset) in WT and more so in all *atg* mutants. Values are means ± SD, *n* = 6; \**P* < 0.05, \*\**P* < 0.01. (B) A typical series of siliques of primary inflorescence in WT and *atg* mutants grown at 23 °C (upper panels), 30 °C (middle panels), and 30 °C after pollination with pollen of the same lines that developed at 23 °C (30 °C × 23 °C: bottom panels). (C) Lengths of siliques in each experiment in B (±SD, *n* = 5). One-way ANOVA followed by Tukey’s *post hoc* test (*P* < 0.05). Bars with the same letter are not significantly different.

We used crossing experiments to investigate whether the HT susceptibility of *atg* mutants was associated with the development of either pollen grains or ovules. When the pistils of *atg* plants that had developed at 30 °C (for 7 days) were pollinated with the pollen of the same lines that had developed at 23 °C, silique length was significantly recovered (Fig. 1B, C), indicating that HT affected pollen fertility but not ovules.

Since autophagy deficiency promotes the activation of SA signaling (Yoshimoto et al. 2009), we monitored the HT sensitivity of WT and *atg5-1* plants carrying a bacterial SA hydroxylase, NahG. The results of Figure 2 and Table 1 show that NahG was unable to rescue the severe sterility of the *atg5-1* mutant.

**Figure 2.**
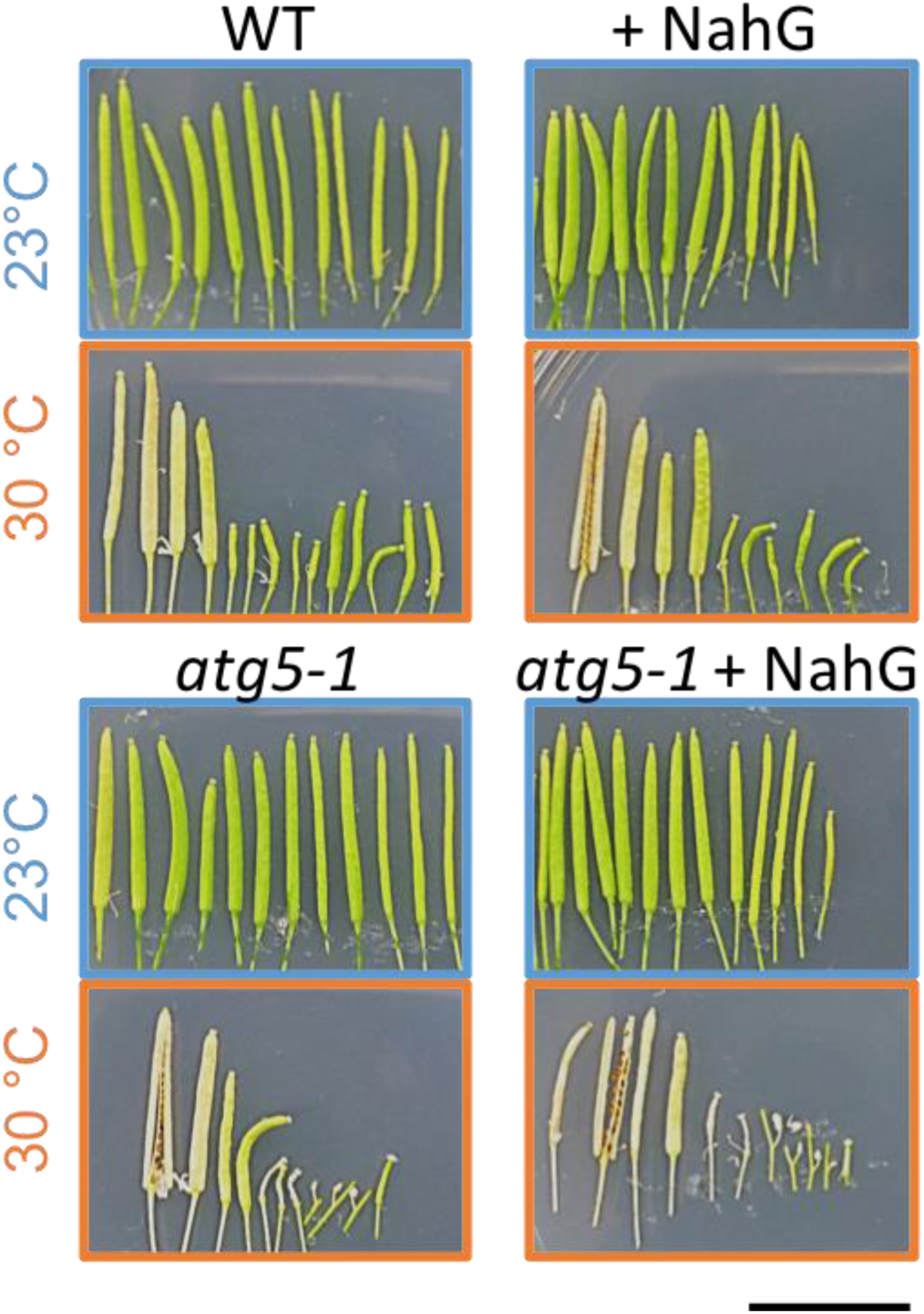
Salicylic acid signaling does not affect hypersensitivity to HT stress in *atg5-1* mutant. Panels show typical series of siliques of primary inflorescence in WT (upper panels) and *atg5-1* plants (lower panels) without (left) or with (right) ectopic NahG to suppress salicylic acid signaling. At 23 °C (blue borders), growth was normal. At 30 °C (orange borders), silique length was drastically reduced from the 5th flower. Scale bars, 1 mm.

We also tested the effect of HT (at 30 and 35 °C) during the seedling stage on root elongation and plant growth. At 30 °C, root growth in both WT and *atg* mutants was retarded by about 20% (Supplementary Fig. 1). At 35 °C, it was retarded by ⅔ or more. Similarly, rosette leaf growth and expansion at 30 °C were suppressed in WT and *atg5-1* (Supplementary Fig. 2). In addition, bolting time was advanced by 1 week in both at 30 °C. These results indicate that the degree of HT-induced obstacles did not change between WT and mutants from the seedling stage to the bolting stage. It suggests that autophagy suppresses HT injury to male reproductive development.

### 3.2. *Deterioration of HT injury to anther and pollen development in* atg *mutants*

At flowering stage, pistil morphology of the 5th to the terminal blossoms was normal in both WT and *atg* mutants, even after transfer to 30 °C (Fig. 3A). In contrast, anther size was reduced by approximately 20 to 40 % at 30 °C in all lines (Fig. 3B). The number of mature pollen grains was decreased in all anthers, severely so in *atg* anthers (Fig. 3B). Anther dehiscence still occurred in WT, even at 30 °C, and pollen grains were released, but anthers in all *atg* mutants barely dehisced (Fig. 3B).

**Figure 3.**
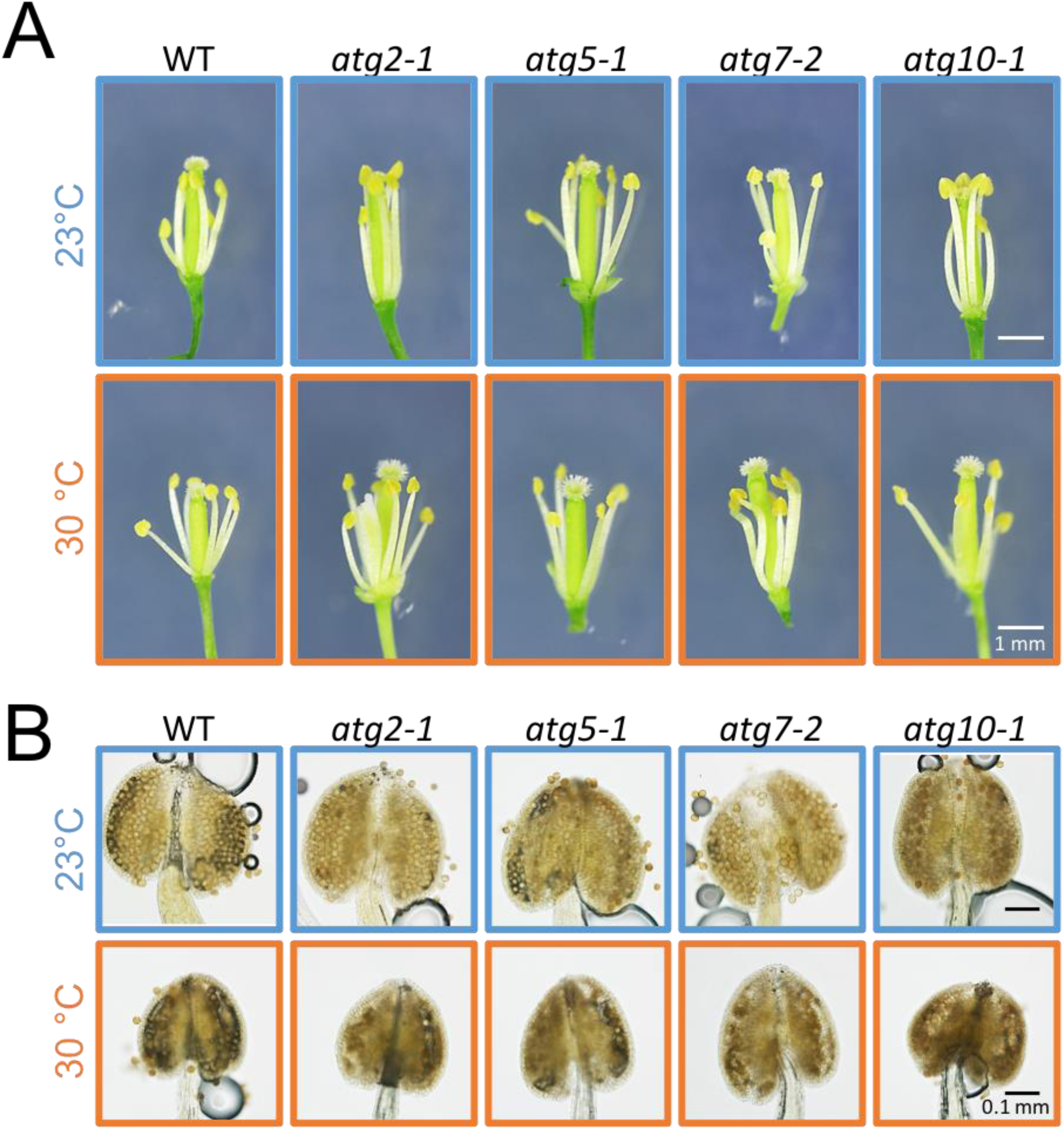
Comparisons of stamen and pistil phenotypes at anthesis. (A) Pistil morphology in WT and *atg* mutants at flowering stage at 23 and 30 °C. Scale bars, 1 mm. (B) Iodine staining of pollen grains in anther of WT and *atg* mutants at 23 and 30 °C. Scale bars, 0.1 mm.

In cross-sections of WT and *atg5-1* stage 13 anthers, at 23 °C, both anthers showed both degeneration of septa in anther locules and breakage along the stomium (Fig. 4A). At 30 °C, WT anthers showed septum and stomium breakage, but *atg5-1* anthers did not (Fig. 4A). In *atg5-1* anthers at 30 °C, some abnormal tapetum cell patches remained, locules shrank, and pollen abortion with larger vacuoles was also frequently observed (Fig. 4A). On the other hand, ovules developed normally in both WT and *atg5-1* under both conditions (Fig. 4B). These results indicate that autophagy is essential for septum and stomium breakage through degeneration of anther wall cells, including in the tapetum, and for pollen maturation at 30 °C.

**Figure 4.**
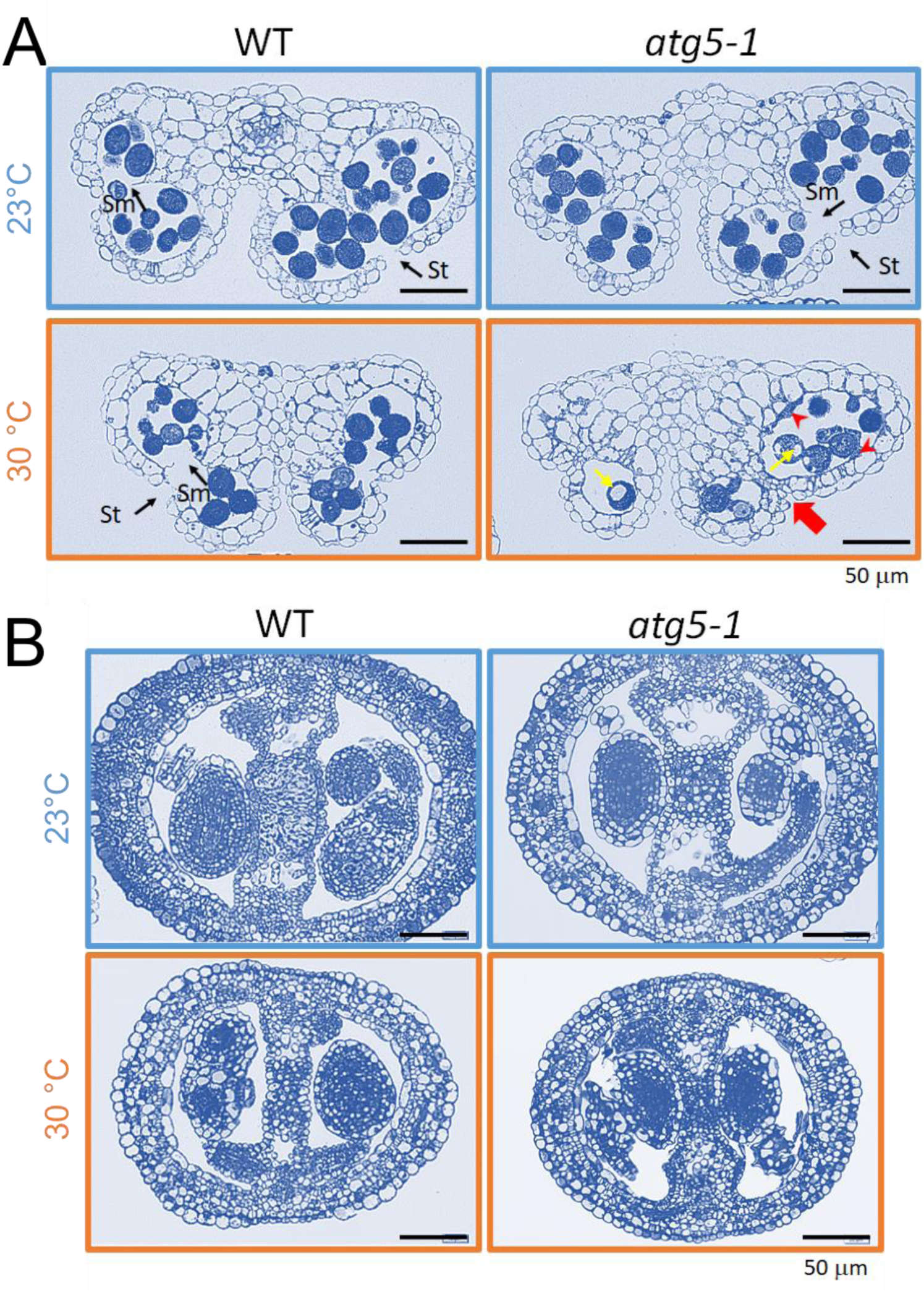
Development of anthers and pistils in WT and *atg5-1* plants at 23 and 30 °C. (A) Transverse sections of stage 13 anthers from flower at anthesis were stained with 0.05% toluidine blue and captured under light microscope. St, stomium; Sm, septum. Scale bars, 50 μm. Red arrowheads, abnormal tapetal patches; red arrow, unbroken septum; yellow arrow, pollen with abnormal large vacuoles. (B) Transverse sections of stage 13 pistils from flower at anthesis.

### 3.3. *Enhanced oxidative stress in anther of* atg5-1 *mutant*

In general, reactive oxygen species (ROS) promote PCD, and spatiotemporal coordination of ROS production is essential for tapetal PCD progression and pollen development in rice and Arabidopsis (Hu et al. 2011; Xie et al. 2014). We monitored H_2_O_2_ levels in stage 12 anthers dissected from WT and *atg5-1* plants held for 3 days at 30 °C. In WT, HT exposure significantly increased DAB staining in pollen grains and anther wall cells (Fig. 5). Intriguingly, at 23 °C, staining was significantly higher in the *atg5-1* anthers than in the WT, and at 30 °C, it further increased in the *atg5-1* pollen grains (Fig. 5).

**Figure 5.**
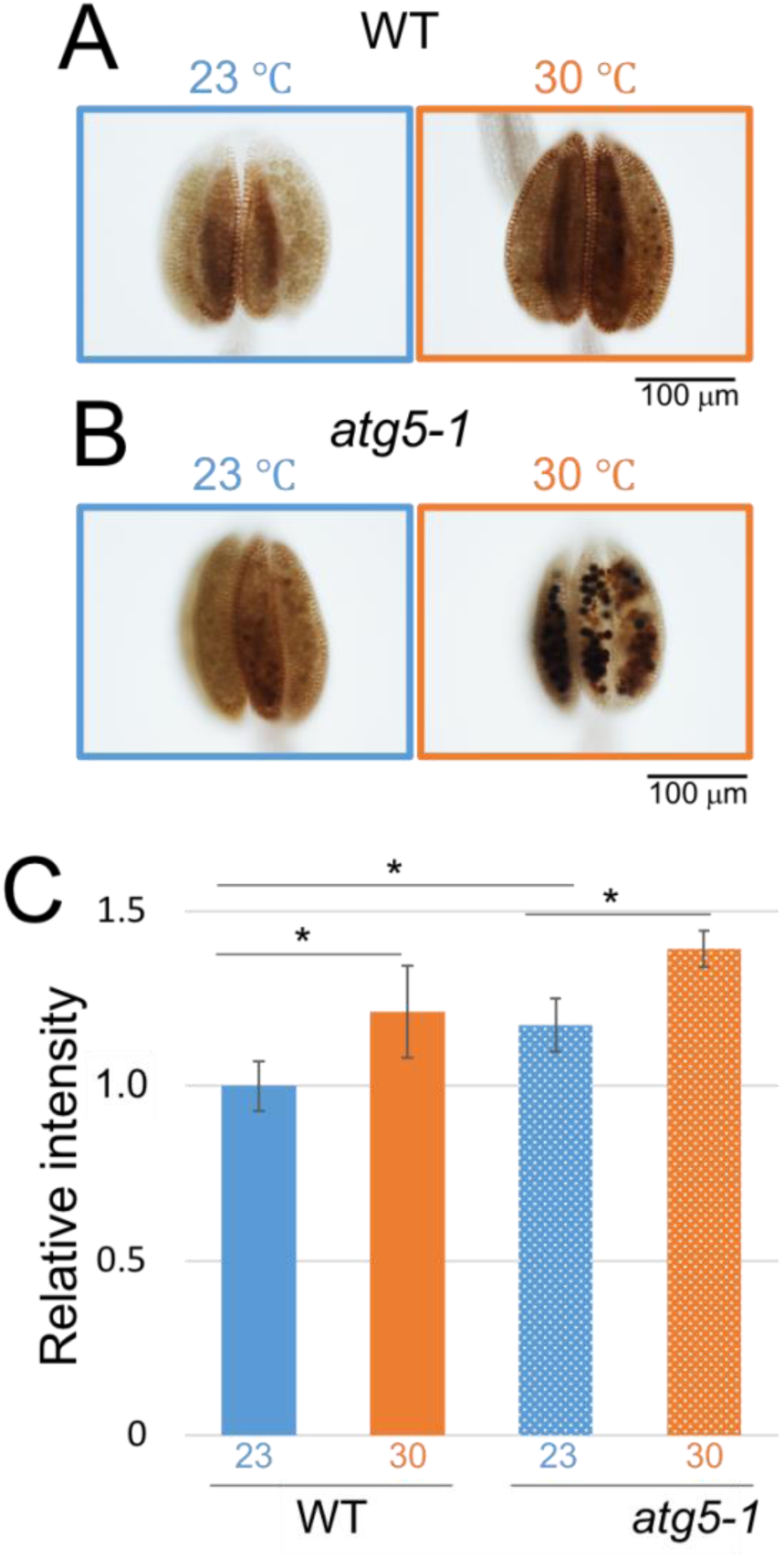
H_2_O_2_ level is increased in stage 12 anthers of *atg5-1* mutant by HT treatment. H_2_O_2_ levels were detected by DAB staining in anthers dissected from (A) WT and (B) *atg5-1* plants after 3 days at 30 °C (orange) or at 23 °C (blue). (C) DAB staining area was quantified by ImageJ software (±SD, *n* = 4). \**P* < 0.05. Scale bars, 100 µm.

SDS-PAGE analysis showed that the intensity of the 24–26-kDa protein band normalized to 41-kDa actin significantly increased in *atg5-1* stage 12 anthers held at 30 °C for 3 days (Fig. 6A, B). To identify which proteins increased, we analyzed the 24–26-kDa bands by MALDI-TOF MS. The major proteins detected were DHAR1, APX1, CTIMC, and 60S ribosomal protein L7-2 (RPL7B). Semi-quantitative MS peak analysis with normalization to RPL7B indicated that HT exposure significantly increased the level of DHAR1, which is involved in scavenging ROS (Dixon et al. 2002), in both WT and *atg5-1* anthers, even though the basal level of DHAR1 at 23 °C was already higher in the mutant (Fig. 6C). Similarly, the basal level of CTIMC, a stress-inducible enzyme of the glycolysis and gluconeogenesis pathway (Sarry et al. 2006), was higher in the *atg5-1* anthers and was induced by HT exposure, but to a lesser extent (Fig. 6C). In both WT and *atg5-1*, HT significantly induced APX1 (Fig. 6C). DHAR1 protein levels were consistent with DAB staining of H_2_O_2_ in WT and *atg5-1* anthers in both control and HT groups.

**Figure 6.**
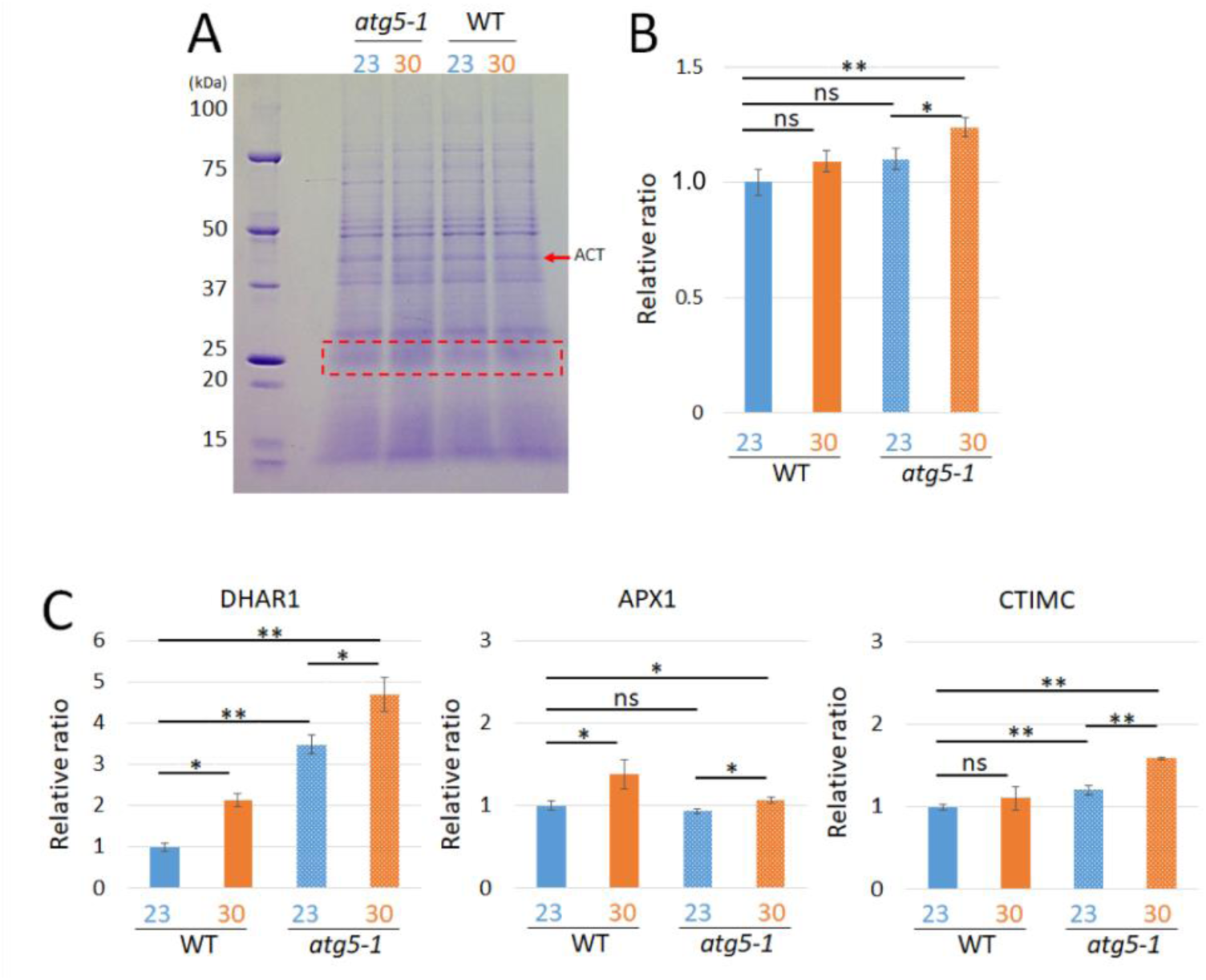
Protein analyses of stage 12 anthers of WT and *atg5-1* plants after HT treatment. (A) Total proteins after 3 days at 30 °C or at 23 °C (control) were separated by 5%–20% gradient SDS PAGE. The proteins at 24–26 kDa (red dashed rectangle) were isolated for MALDI-TOF MS analysis. (B) 24–26-kDa protein band intensity normalized to 41-kDa actin (ACT) (±SD, *n* = 4). (C) DHAR1, APX1, and CTIMC protein levels determined by semi-quantitative MALDI-TOF MS analysis were normalized to 60S ribosomal protein L7-2. \**P* < 0.05, \*\**P* < 0.01.

### 3.4. Autophagy induced in anther wall cells and microspores by moderate HT

To study whether moderate HT induces autophagy in the developing anthers, we investigated the formation of ATG8 foci in *YFP-ATG8e* recombinant plants. GFP-ATG8 fusion proteins are used as reliable molecular markers of autophagy in yeast, mammalian, and Arabidopsis cells (Mizushima et al. 2004; Xie et al. 2008; Ishida et al. 2008; Nakayama et al. 2012; Merkulova et al. 2014). The number of YFP-ATG8e foci significantly increased in the anther wall cells and microspores of stage 9 anthers of plants grown at 30 °C for 3 days (Fig. 7). The foci measured 1–2 µm, comparable to the size of autophagic bodies marked with the GFP-ATG8 fusion protein in Arabidopsis (Li et al. 2014; Lin et al. 2015; Chung et al. 2010).

**Figure 7.**
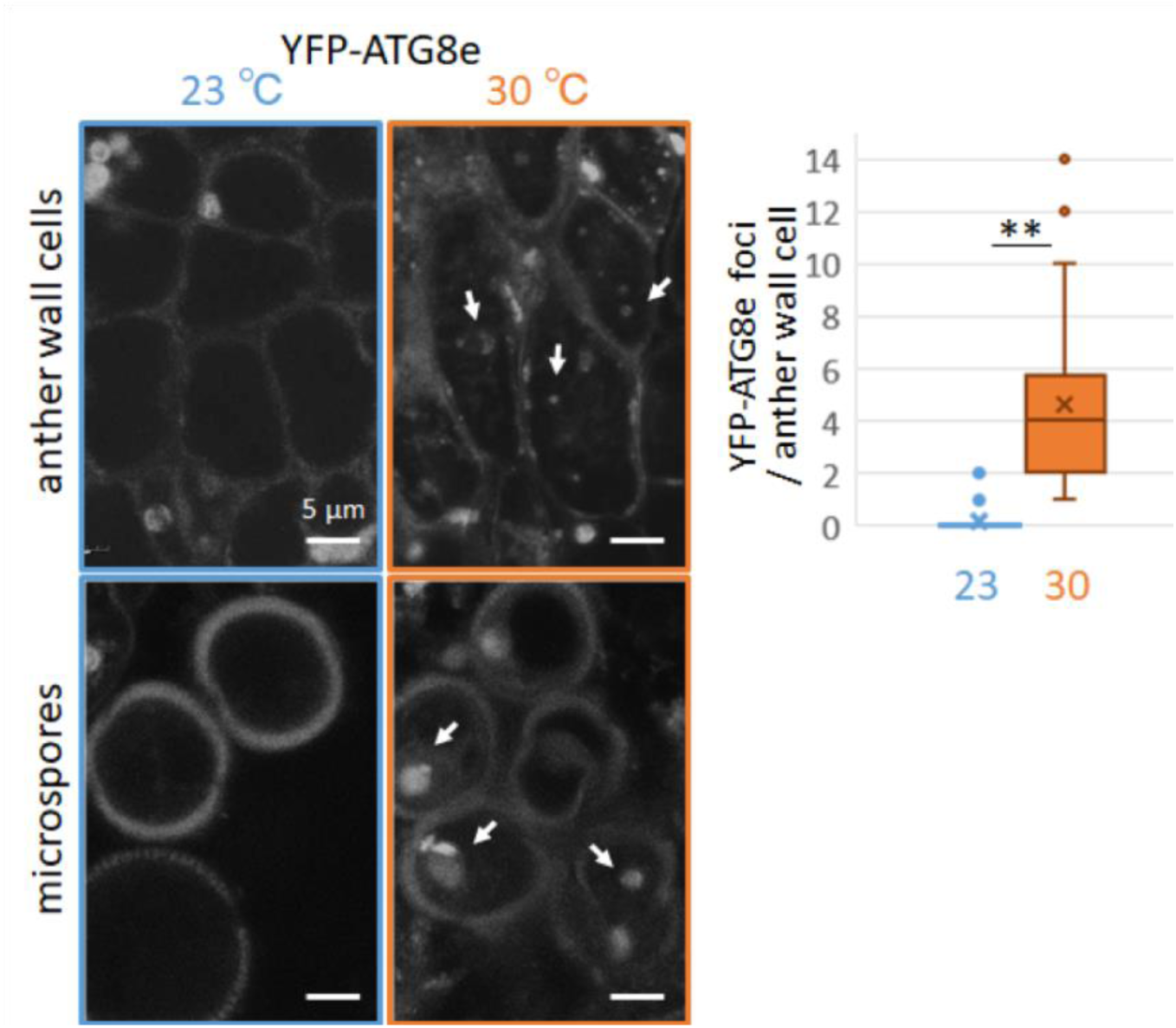
HT stress increased the formation of YFP-ATG8e foci in cells of stage 9 anthers. (Left) Confocal microscopy images of developing anthers dissected from *YFP-ATG8e*-expressing plants grown at 23 °C (blue) or for 3 days at 30 °C (orange). White arrows show YFP-ATG8 foci in anther wall cells (upper) and microspores (lower). (Right) Quantification of YFP-ATG8e foci per anther wall cell (mean ± SD from three independent experiments, >10 cells per sample). \*\**P* < 0.01.

To further investigate how autophagy was promoted, we observed stage 9 anthers of WT and *atg5-1* plants grown at 23 C or for 3 days at 30 °C by transmission electron microscopy. At 23 °C, the tapetosomes and elaioplasts (components of the pollen coat) developed normally in tapetal walls of both WT and *atg5-1* (Fig. 8). A few autophagosomes had the typical double membrane structure in the WT microspores at 23 °C (Fig. 8 inset). At 30 °C, WT tapetosomes developed abnormal vacuoles, and elaioplasts developed larger vacuoles with irregularly fused plastoglobuli (Fig. 8). In addition, the WT microspores showed an increase in autolysosomes, in which lipid bodies were trapped in the vacuoles and gained electron density (Fig. 8). Moreover, the number of autophagosomes increased in WT microspores and anther wall cells (Fig. 8). In contrast, larger vacuoles often appeared in the tapetum cells of *atg5-1* grown at 23 °C (Fig. 8, blue arrows). At 30 °C, the microspores and tapetum cells were shrunken and had increased electron density, but neither vacuolization of elaioplasts and tapetosomes nor lipophage-like structures nor autophagosomes were observed in *atg5-1* (Fig. 8).

**Figure 8.**
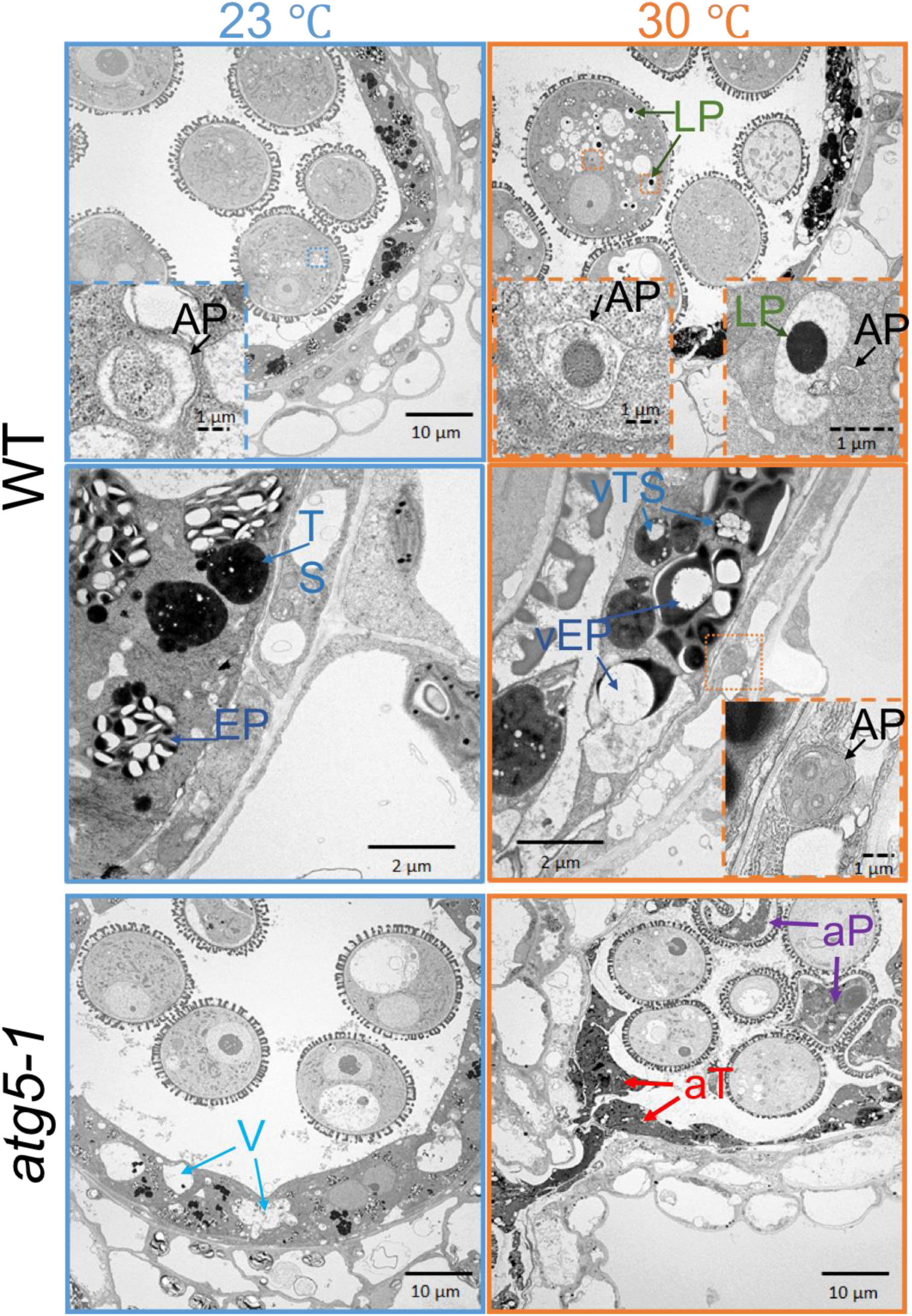
HT induced autophagy in microspores and anther wall cells. Transmission electron microscopy images of stage 9 anther in WT (upper and middle panels) and *atg5-1* mutant (lower panels) at 23 °C or after 3 days at 30 °C. AP: typical autophagosome with double membrane structure; LP: autolysosome including lipid body (lipophage); TS: tapetosome; EP: elaioplast; vTS: tapetosome with abnormal larger vacuoles; vEP: elaioplasts that developed larger vacuoles with irregularly fused plastoglobuli; V: larger vacuoles in *atg5-1* tapetum cells; aT: abnormally shrunken tapetum cells; aP: abnormally shrunken microspores. Scale bars: solid, larger images; dashed, inset images.

### 3.5. *HT repressed transcription of* MYB80 *and* UNDEAD *in developing anthers*

The MYB80 transcription factor is required for the regulation of tapetal PCD; it upregulates *UNDEAD*, which encodes an A1 aspartic protease. It has been suggested that the AtMYB80/ UNDEAD system may regulate the timing of tapetal PCD, which is critical for viable pollen production (Phan et al. 2011, 2012). To elucidate whether HT affects MYB80 transcriptional regulation in developing anther cells, we analyzed the expression of *MYB80* and *UNDEAD* in stage 9 anthers of WT and *atg5-1* plants. HT treatment at 30 °C for 3 days significantly reduced the expression of both *MYB80* and *UNDEAD* in both WT and *atg5-1* relative to 23 °C (Fig. 9). In particular, *UNDEAD* expression was much lower in *atg5-1* than in WT at 30 °C, suggesting that normal tapetal PCD is considerably inhibited in *atg5-1* under HT.

**Figure 9.**
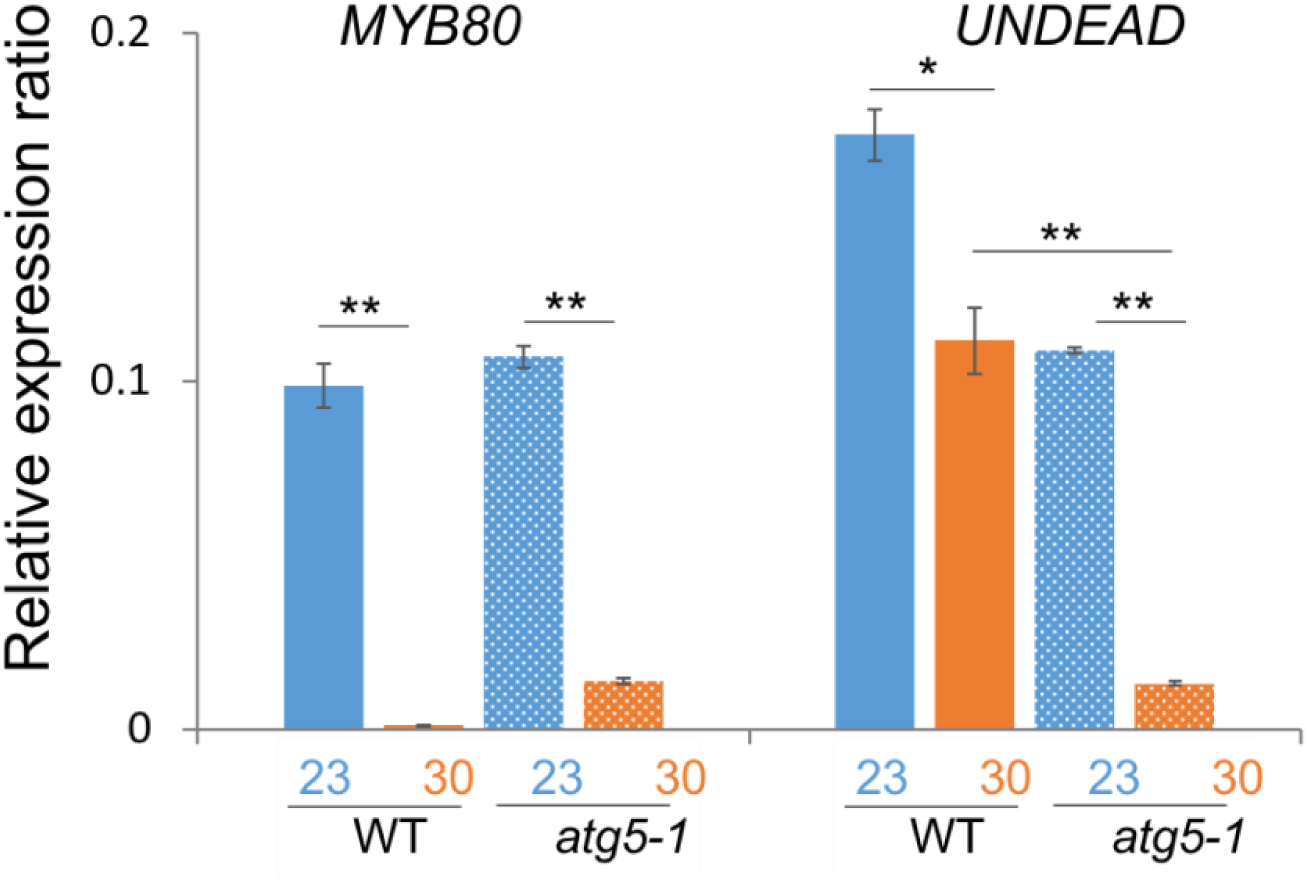
Altered expression of MYB80 signaling genes *MYB80* and *UNDEAD* in stage 9 anthers of WT and *atg5-1* plants grown at 23 °C or after 3 days at 30 °C. Expression levels were monitored using real-time quantitative PCR and normalized to expression of *Tubulin 2/3*. Values are means ± SD. \**P* < 0.05; \*\**P* < 0.01.

## 4. Discussion

Plant HT injury is one of the major obstacles to crop yield under warmer temperatures (Lobell and Field 2007). We analyzed the role of autophagy in the effect of HT stress on reproductive development of Arabidopsis. Although the roles of autophagy in senescence and in responses to nutrient starvation, pathogens, drought, salt, and oxidative stresses are well known (Doelling et al. 2002; Hanaoka et al. 2002; Bassham et al. 2006; Chung et al. 2009; Hofius et al. 2011; Yoshimoto 2012), there are few reports of its role in HT stress. Zhou et al. (2014) reported that silencing of *ATG*-related genes reduced the tolerance of tomato to severe heat stress at 45 °C. Although HT stress affects the whole plant, microsporogenesis is one of the most sensitive processes (Sakata & Higashitani 2008; Müller & Rieu 2016).

We previously showed that HT increases vacuolization and the development of autolysosome-or autophagosome-like structures in developing anther cells of barley (Oshino et al. 2007, 2011). Singh et al. (2010, 2015) showed that overexpression of the Arabidopsis tapetum-cell-specific autophagy-related *ATG6/BECLIN1* in tobacco leads to premature degeneration of the developing tapetum cells. These reports suggest that disrupting autophagy would prevent the premature degradation of tapetum cells and improve HT tolerance. Instead, our results show that autophagy is essential for mitigating HT injury during pollen development in Arabidopsis: *atg* mutants became almost completely male-sterile at moderate HT of 30 °C, whereas WT plants still produced some fertile seeds (Figs. 1, 2; Table 1). In all *atg* mutants tested, 30 °C inhibited anther dehiscence and pollen development, leading to male sterility (Figs. 2, 3). Transverse section analysis of *atg5-1* anthers clearly revealed the inhibition of anther septum and stomium breakage and shrunken abnormal tapetum (Fig. 4).

Anther development and differentiation, including specification of cell lineage and cell fate, are well regulated programs. The epidermis, endothecium, middle layer, and tapetum of anther wall cells are sequentially degraded during pollen maturation and anther dehiscence. The proper timing of tapetal degradation is necessary for the production of viable pollen, and occurs via developmentally regulated PCD (Papini et al. 1999; Varnier et al. 2005; Parish & Li 2010). MYB80 is a key transcription factor that controls tapetal PCD by regulating at least 70 genes, including directly upregulating *UNDEAD* (Phan et al. 2011, 2012; Xu et al. 2014). In our study, *MYB80* expression in WT and *atg5-1* anthers decreased similarly at 30 °C (Fig. 9), so increasing temperatures inhibited tapetal PCD.

ATG8e foci increased in tapetum cells and microspores at 30 °C (Fig. 7). Size and number of vacuoles increased in the cytoplasm, elaioplasts, and tapetosomes, and autophagosomes, autolysosomes, and lipophage-like structures increased in microspores and anther wall cells (Fig. 8). In mammalian cells, the breakdown of lipid droplets by lipophagy contributes to energy generation under stress (Dong & Czaja 2011). In the *atg5-1* mutant, these autophagic phenomena did not appear at 30 °C. Instead, abnormal large vacuolization and subsequent shrinkage (but not complete degeneration) of tapetum cells were observed, and the collapse of microspores resulted in complete male sterility (Figs. 4, 8). Together, these results indicate that HT stress alters the timing of tapetal PCD through repression of MYB80 transcriptional regulation, but it activates autophagy to compensate for the alteration of MYB80-regulated PCD by autophagic cell death for rescuing microspore development under HT.

A rice autophagy-defective mutant, the OsATG7 Tos 17 insertion line, is male-sterile owing to the inhibition of tapetum-cell degradation (Kurusu et al. 2014; Hanamata et al. 2014). Autophagy is involved in the breakdown of the lipid bodies and of lipids transferred from tapetum cells to the microspore surface in rice (Kurusu et al. 2014; Hanamata et al. 2014). However, Arabidopsis *atg* mutants completed their life cycle at normal temperature (Yoshimoto 2012; Figs. 1–4, Supplementary Figs. 1, 2), and thus rice and Arabidopsis differ in their need for autophagy (Kurusu et al. 2014; Hanamata et al. 2014). In brassicaceous species, including Arabidopsis, a distinctive organelle, the tapetosome, develops in the tapetum cells from the endoplasmic reticulum and stores triacylglycerols, flavonoids, alkanes, and pollen coat proteins (Wu et al. 1997; Suzuki et al. 2013; Hanamata et al. 2014). Another tapetal organelle, the elaioplast, stores a major component of the pollen coat: steryl esters inside numerous plastoglobuli (Wu et al. 1997; Suzuki et al. 2013; Hanamata et al. 2014). Under HT, where tapetal PCD activity decreased, we found lots of vacuoles in tapetosomes and elaioplasts in WT anther wall cells, and lipophage-like bodies trapped in the autophagosomes and vacuoles in microspores (Fig. 8), similar to previous observations in rice (Kurusu et al. 2014; Hanamata et al. 2014), but not in *atg5-1* cells (Fig. 8). These results strongly suggest that degradation of elaioplasts and tapetosomes promoted by autophagy leads to degeneration of tapetum cells and degrades lipid bodies in microspores to prevent abortion of microspores by inhibiting PCD at elevated temperatures.

In addition to the hypothesis that autophagic cell death can compensate for the decrease of tapetal PCD by HT, the following hypothesis is also conceivable. Here we found that HT induces ROS in developing anthers of both *atg5-1* and WT plants (Fig. 5). Both APX1 and DHAR1 levels were significantly induced following HT exposure, DHAR1 greatly so (Fig. 6C). Recently, a similar pattern of APX1 and DHAR1 upregulation was reported in salt-stressed Arabidopsis roots (Szymanska et al. 2019). Moreover, even under steady-state conditions, H_2_O_2_ and DHAR1 levels were higher in developing anthers of *atg5-1* than in WT (Figs. 5, 6). Likewise, higher ROS in *atg2-1* and *atg5-1* leaves was reported (Yoshimoto et al. 2009). In addition, it has recently been shown that excess ROS generated in a cytoplasmic-male-sterile wheat line delayed tapetal PCD initiation and led to pollen abortion (Liu et al. 2018), although the spatiotemporal regulation of ROS production is essential for tapetal PCD progression (Hu et al. 2011; Xie et al. 2014). Altogether, in addition to HT-induced ROS, over-accumulation of ROS in autophagy deficiency might strongly affect the spatiotemporal ROS signaling for tapetal PCD. In either hypothesis, the proper timing of pollen development, including tapetal PCD, is strictly controlled, and autophagy is important to minimize the effects of HT and to maintain this pollen-specific development.

## 5. Conclusion

In Arabidopsis, autophagy is not essential for completion of the life cycle under normal temperatures. Seedling growth of autophagy-deficient mutants was unaffected by HT. However, pollen development was significantly damaged by HT: HT stress induced autophagy in developing anther wall cells and microspores. It caused oxidative damage and altered the tapetal PCD, but activated autophagy mitigated the HT injury.

## Author contributions

AH and GD designed all experiments; GD, MI and AH wrote the manuscript; ZS, GD, NH, MK, SN, and MI performed experiments and analyzed the data. All authors approved the manuscript.

## Acknowledgements

We thank Dr. H. Ishida, Tohoku University (Sendai, Japan), Dr. K. Yoshimoto, Meiji University (Tokyo, Japan), and the Nottingham Arabidopsis Resource Centre (Nottingham, UK) for kindly supplying the *atg* mutants and GFP recombinant lines; Dr. H. Ishida and Dr. K. Kuchitsu, Tokyo University of Science (Chiba, Japan) for helpful suggestions; Dr T. Nakagawa, Shimane University (Matsue, Japan), for providing the R4pGWB vectors; and Dr. Christopher Grefen, University of Tübingen (Tübingen, Germany), for providing the pUBN vectors. This work was funded in part by the Ministry of Education, Culture, Sports, Science and Technology grants 15H04616 and 26506001.

---

**Supplementary Fig. 1** Effect of HT (30 and 35 °C) on seedling root growth in WT and *atg* mutant plants. (Upper) Root growth of 4-day-old seedlings at 23, 30, or 35 °C on 1/2 MS agar. (Lower) Daily root length (±SD, *n* = 4).

**Supplementary Fig 2**. HT reduces vegetative growth and accelerates flowering in both WT and *atg5-1* mutant. Images show plants grown in soil for (A) 10 days at 23 or 30 °C and (B) 30 days at 23 °C or 22 days at 30 °C. Scale bars, 1 cm.

## References

Abiko M, Akibayashi K, Sakata T, Kimura M, Kihara M, Itoh K, Asamizu E, Sato S, Takahashi H, Higashitani A. (2005) High-temperature induction of male sterility during barley (*Hordeum vulgare* L.) anther development is mediated by transcriptional inhibition. Sex Plant Reprod 18: 91–100. DOI: 10.1007/s00497-005-0004-2

Baehrecke EH (2005) Autophagy: dual roles in life and death? Nat Rev Mol Cell Biol 6: 505–510. DOI: 10.1038/nrm1666

Bassham DC, Laporte M, Marty F, Moriyasu Y, Ohsumi Y, Olsen LJ, Yoshimoto K. (2006) Autophagy in development and stress responses of plants. Autophagy 2: 2–11. DOI: 10.4161/auto.2092

Chung T, Suttangkakul A, Vierstra RD. (2009) The ATG autophagic conjugation system in maize: ATG transcripts and abundance of the ATG8–lipid adduct are regulated by development and nutrient availability. Plant Physiol 149: 220–234. DOI: 10.1104/pp.108.126714

Chung T, Allison R. Phillips and Richard D. Vierstra (2010) ATG8 lipidation and ATG8-mediated autophagy in Arabidopsis require ATG12 expressed from the differentially controlled *ATG12A* and *ATG12B* loci. Plant J (2010) 62, 483–493. doi: 10.1111/j.1365-313X.2010.04166

Clough SJ, Bent AF. (1998) Floral dip: simplified method for *Agrobacterium*-mediated transformation of *Arabidopsis thaliana*. Plant J 16: 735–743. DOI: 10.1046/j.1365-313x.1998.00343.X

Dixon DP, Davis BG, Edwards R. (2002) Functional divergence in the glutathione transferase superfamily in plants. Identification of two classes with putative functions in redox homeostasis in Arabidopsis thaliana. J Biol Chem. 277: 30859–30869. DOI: 10.1074/jbc.M202919200

Doelling JH, Walker JM, Friedman EM, Thompson AR, Vierstra RD. (2002) The APG8/12-activating enzyme APG7 is required for proper nutrient recycling and senescence in *Arabidopsis thaliana*. J Biol Chem 277: 33105–33114. DOI: 10.1074/jbc.M204630200

Dong H, Czaja M J. (2011) Regulation of lipid droplets by autophagy [J]. Trends Endocr Metab 22: 234–240. DOI: 10.1016/j.tem.2011.02.003

Grefen C, Donald N, Hashimoto K, Kudla J, Schumacher K, Blatt MR. (2010) A ubiquitin-10 promoter-based vector set for fluorescent protein tagging facilitates temporal stability and native protein distribution in transient and stable expression studies. Plant J 64: 355–365. DOI: 10.1111/j.1365-313X.2010.04322.X.

Hanamata S, Kurusu T, Kuchitsu K. (2014) Roles of autophagy in male reproductive development in plants. Front Plant Sci 5: 457. DOI: 10.3389/fpls.2014.00457.

Hanaoka H, Noda T, Shirano Y, Kato T, Hayashi H, Shibata D, Tabata S, Ohsumi Y. (2002) Leaf senescence and starvation-induced chlorosis are accelerated by the disruption of an Arabidopsis autophagy gene. Plant Physiol 129: 1181–1193. DOI: 10.1104/pp.011024

Hofius D, Munch D, Bressendorff S, Mundy J, Petersen M. (2011) Role of autophagy in disease resistance and hypersensitive response-associated cell death. Cell Death Differ 18: 1257–1262. DOI: 10.1038/cdd.2011.43.

Hu L, Liang W, Yin C, Cui X, Zong J, Wang X, Hu J, Zhang D. (2011) Rice MADS3 regulates ROS homeostasis during late anther development. Plant Cell 23: 515–533. DOI: 10.1105/tpc.110.074369.

Ishida H, Yoshimoto K, Izumi M, Reisen D, Yano Y, Makino A, Ohsumi Y, Hanson MR., Mae T. (2008) Mobilization of rubisco and stromatolites localized fluorescent proteins of chloroplasts to the vacuole by an *ATG* gene-dependent autophagic process. Plant Physiol 148: 142–155. DOI: 10.1104/pp.108.122770

Kim I, Rodriguez-Enriquez S, Lemasters JJ (2007) Selective degradation of mitochondria by mitophagy. Arch Biochem Biophys 462: 245–253. DOI: 10.1016/j.abb.2007.03.034

Klionsky DJ, Emr SD (2000) Autophagy as a regulated pathway of cellular degradation. Science 290: 1717–1721. DOI: 10.1126/science.290.5497.1717

Kroemer G, Jäättelä M (2005) Lysosomes and autophagy in cell death control. Nat Rev Cancer 5: 886–897. DOI: 10.1038/nrc1738

Kurusu T, Koyano T, Hanamata S, Kubo T, Noguchi Y, Yagi C, Nagata N, Yamamoto T, Ohnishi T, Okazaki Y, Kitahata N, Ando D, Ishikawa M, Wada S, Miyao A, Hirochika H, Shimada H, Makino A, Saito K, Ishida H, Kinoshita T, Kurata N, Kuchitsu K. (2014) OsATG7 is required for autophagy-dependent lipid metabolism in rice postmeiotic anther development. Autophagy 10: 878–888. DOI: 10.4161/auto.28279.

Li FQ, Taijoon Chung, Richard D. Vierstra. (2014) AUTOPHAGY-RELATED11 plays a critical role in general autophagy-and senescence-induced mitophagy in *Arabidopsis*. The Plant Cell DOI: https://doi.org/10.1105/tpc.113.120014

Lin YS, Yu Ding, Juan Wang, Jinbo Shen, Chun Hong Kung, Xiaohong Zhuang, Yong Cui, Zhao Yin, Yiji Xia, Hongxuan Lin, David G. Robinson, Liwen Jiang (2015). Exocyst-positive organelles and autophagosomes are distinct organelles in plants. Plant Physiol DOI: https://doi.org/10.1104/pp.15.00953

Liu Z, Shi X, Li S, Hu G, Zhang L, Song X. (2018) Tapetal-delayed programmed cell death (PCD) and oxidative stress-induced male sterility of *Aegilops uniaristata* cytoplasm in wheat. Int J Mol Sci 19: pii: E1708. DOI: 10.3390/ijms19061708.

Lobell DB, Field CB. (2007) Global scale climate-crop yield relationships and the impact of recent warming. Environ Res Lett 2: 014002. DOI:10.1088/1748-9326/2/1/014002

Love AJ, Milner JJ, Sadanandom A. (2008) Timing is everything: regulatory overlap in plant cell death. Trends Plant Sci 13: 589–595. DOI: 10.1016/j.tplants.2008.08.006

Merkulova EA, Guiboileau A, Naya L, Daubresse CM, Yoshimoto K (2014) Assessment and optimization of autophagy monitoring methods in Arabidopsis roots indicate direct fusion of autophagosomes with vacuoles. Plant Cell Physiol 55: 715–726 DOI: 10.1093/pcp/pcu041

Mizushima N, Yamamoto A, Matsui M, Yoshimori T, Ohsumi Y (2004) *In vivo* analysis of autophagy in response to nutrient starvation using transgenic mice expressing a fluorescent autophagosome marker. Mol Biol Cell 15: 1101–1111. DOI: 10.1091/mbc.E03-09-0704

Müller F, Rieu I. (2016) Acclimation to high temperature during pollen development. Plant Reprod 29: 107–118. DOI: 10.1007/s00497-016-0282-X

Nakagawa T, Nakamura S, Tanaka K, Kawamukai M, Suzuki T, Nakamura K, Kimura T, Ishiguro S. (2008) Development of R4 gateway binary vectors (R4pGWB) enabling high-throughput promoter swapping for plant research. Biosci Biotechnol Biochem 72: 624–629. DOI: 10.1271/bbb.70678

Nakamura S, Hidema J, Sakamoto W, Ishida H, Izumi M (2018) Selective elimination of membrane-damaged chloroplasts via microautophagy. Plant Physiol 177: 1007–1026. DOI: 10.1104/pp.18.00444

Nakatogawa H, Suzuki K, Kamada Y, Ohsumi Y (2009) Dynamics and diversity in autophagy mechanisms: lessons from yeast. Nat Rev Mol Cell Biol. 10: 458–467. DOI: 10.1038/nrm2708.

Nakayama M, Kaneko Y, Miyazawa Y, Fujii N, Higashitani N, Wada S, Ishida H, Yoshimoto K, Shirasu K, Yamada K, Nishimura M, Takahashi H. (2012) A possible involvement of autophagy in amyloplast degradation in columella cells during hydrotropic response of Arabidopsis roots [J]. Planta 236: 999–1012. DOI: 10.1007/s00425-012-1655-5

Ohsumi Y (1999) Molecular mechanism of autophagy in yeast, *Saccharomyces cerevisiae*. Phil Trans R Soc Lond B Biol Sci 354: 1577–1581. DOI: 10.1098/rstb.1999.0501

Orozco-Cardenas M, Ryan CA (1999) Hydrogen peroxide is generated systemically in plant leaves by wounding and systemin via the octadecanoid pathway. Proc Natl Acad Sci U S A 11: 6553–6557. DOI 96.11.6553

Oshino T, Abiko M, Saito R, Ichiishi E, Endo M, Kawagishi-Kobayashi M, Higashitani A. (2007) Premature progression of anther early developmental programs accompanied by comprehensive alterations in transcription during high-temperature injury in barley plants. Mol Genet Genom 278: 31–42. DOI: 10.1007/s00438-007-0229-X

Oshino T, Miura S, Kikuchi S, Hamada K, Yano K, Watanabe M, Higashitani A. (2011) Auxin depletion in barley plants under high-temperature conditions represses DNA proliferation in organelles and nuclei via transcriptional alterations. Plant Cell Environ 34: 284–290. DOI: 10.1111/j.1365-3040.2010.02242.X

Papini A, Mosti S, Brighigna L. (1999) Programmed-cell-death events during tapetum development of angiosperms. Protoplasma 207: 213–221. DOI: 10.1007/BF01283002

Parish RW, Li SF. (2010) Death of a tapetum: A programme of developmental altruism. Plant Science 178:73–89. DOI: 10.1016/j.plantsci.2009.11.001

Phan HA, Iacuone S, Li SF, Parish RW. (2011) The MYB80 transcription factor is required for pollen development and the regulation of tapetal programmed cell death in *Arabidopsis thaliana*. Plant Cell 23: 2209–2224. DOI: 10.1105/tpc.110.082651

Phan HA, Li SF, Parish RW. (2012) MYB80, a regulator of tapetal and pollen development, is functionally conserved in crops. Plant Mol Biol 78: 171–183. DOI: 10.1007/s11103-011-9855-0

Sakata T, Higashitani A. (2008) Male sterility accompanied with abnormal anther development in plants – Genes and environmental stresses with special reference to high temperature injury. Intl J Plant Dev Biol 2: 42–51.

Sakata T, Oshino T, Miura S, Tomabechi M, Tsunaga Y, Higashitani N, Miyazawa Y, Takahashi H, Watanabe M, Higashitani A. (2010) Auxins reverse plant male sterility caused by high temperatures. Proc Natl Acad Sci U S A 107: 8569–8574. DOI: 10.1073/pnas.1000869107.

Sanders PM, Bui AQ, Weterings K, McIntire KN, Hsu Y-C, Lee PY, Truong MT, Beals TP, Goldberg RB. (1999) Anther developmental defects in *Arabidopsis thaliana* male-sterile mutants. Sex Plant Reprod 11: 297–322.

Sarry JE, Kuhn L, Ducruix C, Lafaye A, Junot C, Hugouvieux V, Jourdain A, Bastien O, Fievet JB, Vailhen D, Amekraz B, Moulin C, Ezan E, Garin J, Bourguignon J. (2006) The early responses of *Arabidopsis thaliana* cells to cadmium exposure explored by protein and metabolite profiling analyses. Proteomics 6: 2180–2198. DOI: 10.1002/pmic.200500543

Sasagawa Y, Kikuchi K, Dazai K, Higashitani A. (2005) Caenorhabditis elegans Elongin BC complex is essential for cell proliferation and chromosome condensation and segregation during mitosis and meiotic division II. Chromosome Res 13: 357–375. DOI: 10.1007/s10577-005-2687-5

Singh SP, Pandey T, Srivastava R, Verma PC, Singh PK, Tuli R, Sawant SV. (2010) BECLIN1 from Arabidopsis thaliana under the generic control of regulated expression systems, a strategy for developing male sterile plants. Plant Biotechnol J 8: 1005–1022. DOI: 10.1111/j.1467-7652.2010.00527.X

Singh SP, Singh SP, Pandey T, Singh RR, Sawant SV. (2015) A novel male sterility-fertility restoration system in plants for hybrid seed production. Sci Rep 5: 11274. DOI: 10.1038/srep11274

Smyth DR, Bowman JL, Meyerowitz EM. (1990) Early flower development in Arabidopsis. Plant Cell 2: 755–767. DOI: 10.1105/tpc.2.8.755

Suzuki T, Tsunekawa S, Koizuka C, Yamamoto K, Imamura J, Nakamura K, Ishiguro S. (2013) Development and disintegration of tapetum-specific lipid-accumulating organelles, elaioplasts and tapetosomes, in *Arabidopsis thaliana* and *Brassica napus*. Plant Sci 207: 25–36. DOI: 10.1016/j.plantsci.2013.02.008.

Szymanska KP, Kowalczyk LP, Lichocka M, Maszkowska J and Dobrowolska G (2019) SNF1-Related Protein Kinases SnRK2.4 and SnRK2.10 Modulate ROS Homeostasis in Plant Response to Salt Stress. Int J Mol Sci 20: 143; DOI:10.3390/ijms20010143

Thompson, AR, Doelling JH, Suttangkakul A, Vierstra RD (2005) Autophagic nutrient recycling in Arabidopsis directed by the ATG8 and ATG12 conjugation pathways. Plant Physiol 138: 2097–2110

Varnier AL, Mazeyrat-Gourbeyre F, Sangwan RS, Clement C (2005) Programmed cell death progressively models the development of anther sporophytic tissues from the tapetum and is triggered in pollen grains during maturation. J Struct Biol 152:118–128. DOI: 10.1016/j.jsb.2005.07.011

Wada S, Hayashida Y, Izumi M, Kurusu T, Hanamata S, Kanno K, Kojima S, Yamaya T, Kuchitsu K, Makino A, Ishida H. (2015) Autophagy supports biomass production and nitrogen use efficiency at the vegetative stage in rice. Plant Physiol 168: 60–73. DOI: 10.1104/pp.15.00242.

Wu SS, Platt KA, Ratnayake C, Wang TW, Ting JT, Huang AH. (1997) Isolation and characterization of neutral-lipid-containing organelles and globuli-filled plastids from Brassica napus tapetum. Proc Natl Acad Sci U S A 94: 12711–12716. DOI: 10.1073/pnas.94.23.12711

Xia K, Liu T, Ouyang J, Wang R, Fan T, Zhang M. (2011) Genome-wide identification, classification, and expression analysis of autophagy-associated gene homologues in rice (*Oryza sativa* L.). DNA Res 18: 363–377. DOI: 10.1093/dnares/dsr024

Xie H-T, Wan Z-Y, Li S, Zhang Y. (2014) Spatiotemporal production of reactive oxygen species by NADPH oxidase is critical for tapetal programmed cell death and pollen development in Arabidopsis. Plant Cell 26: 2007–2023. DOI: 10.1105/tpc.114.125427

Xie Z, Nair U, Klionsky DJ. (2008) Atg8 controls phagophore expansion during autophagosome formation. Mol Biol Cell 19: 3290–3298. DOI: 10.1091/mbc.E07-12-1292

Xu Y, Iacuone S, Li SF, Parish RW. (2014) MYB80 homologues in Arabidopsis, cotton and Brassica: regulation and functional conservation in tapetal and pollen development. BMC Plant Biol 14: 278. DOI: 10.1186/s12870-014-0278-3

Yorimitsu T, Klionsky DJ (2005) Autophagy: Molecular machinery for self-eating. Cell Death Differ 12: 1542–1552. DOI: 10.1038/sj.cdd.4401765

Yoshimoto K. (2012) Beginning to understand autophagy, an intracellular self-degradation system in plants. Plant Cell Physiol 53: 1355–1365. DOI: 10.1093/pcp/pcs099

Yoshimoto K. Jikumaru Y, Kamiya Y, Kusano M, Consonni C, Panstruga R, Ohsumi Y, Shirasu K. (2009) Autophagy negatively regulates cell death by controlling NPR1-dependent salicylic acid signaling during senescence and the innate immune response in Arabidopsis. Plant Cell 21: 2914–2927. DOI: 10.1105/tpc.109.068635

Zhou J. Wang J. Yu J-Q, Chen Z. (2014) Role and regulation of autophagy in heat stress responses of tomato plants. Front Plant Sci 5: 174. DOI: 10.3389/fpls.2014.00174

