## Supplementary figures and images for "Autophagy Mitigates High-Temperature Injury in Pollen Development of *Arabidopsis thaliana*"

### Supplementary Fig. 1, Supplementary Fig. 2

#### Slide 1
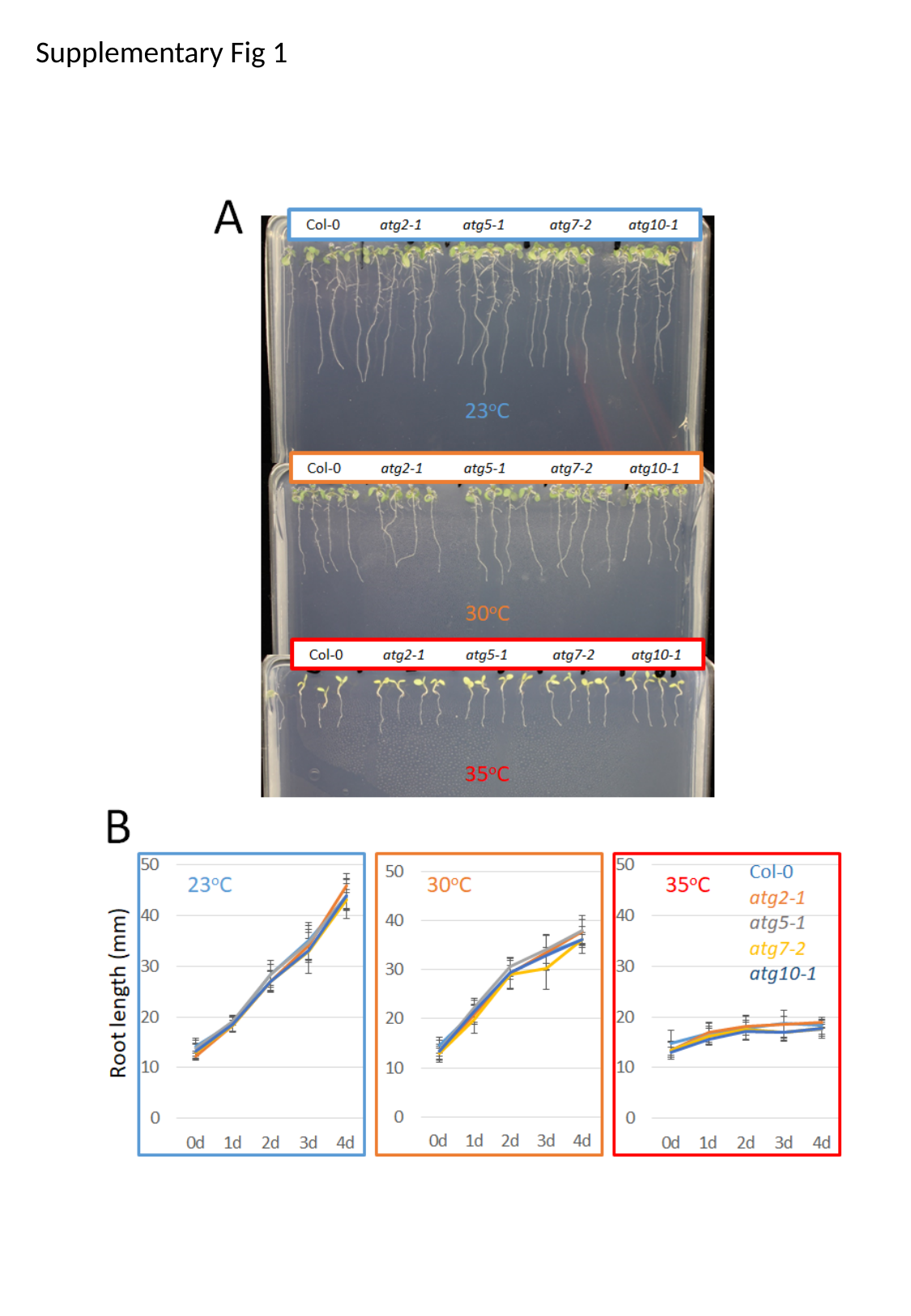

Supplementary Fig 1

#### Slide 2
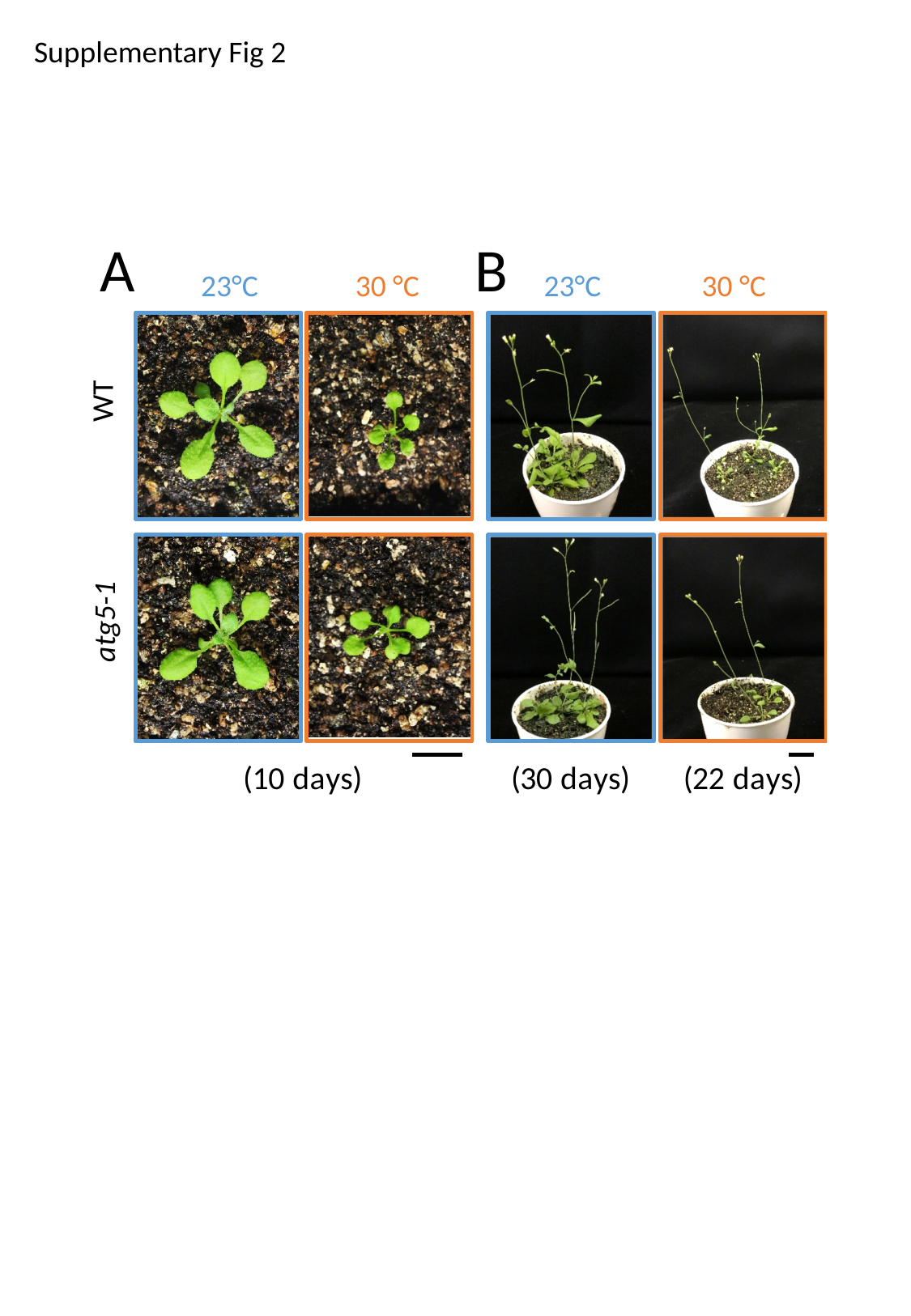

Supplementary Fig 2
A
B
23°C
30 °C
23°C
30 °C
WT
atg5-1
